# *BioSankey*: Visualizing microbial communities and gene expression data over time

**DOI:** 10.1101/191767

**Authors:** Alexander Platzer, Julia Polzin, Ping Penny Han, Klaus Rembart, Thomas Nussbaumer

## Abstract

Metagenomics, RNA-seq, WGS (Whole Genome Sequencing) and other types of next-generation sequencing techniques provide quantitative measurements for single strains and genes over time. To obtain a global overview of the experiment and to explore the full potential of a given dataset, intuitive and interactive visualization tools are needed. Therefore, we established *BioSankey*, which allows to visualize microbial species in microbiome studies and gene expression over time as a Sankey diagram. These diagrams are embedded into a project-specific HTML page, that contains all information as provided during the installation process. *BioSankey* can be easily applied to analyse bacterial communities in time-series datasets. Furthermore, it can be used to analyse the fluctuations of differentially expressed genes (DEG). The output of *BioSankey* is a project-specific HTML page, which depends only on JavaScript to enable searches of interesting species or genes of interest without requiring a web server or connection to a database to exchange results among collaboration partners. *BioSankey* is a tool to visualize different data elements from single and dual RNA-seq datasets as well as from metagenomes studies.

## Introduction

Dramatic reductions in sequencing costs and improvements of sequencing technologies have led to a higher throughput of sequence material in much shorter time and led to a burst of dual transcriptome experiments and metagenome studies over the last years. To understand and to find key genes in these datasets, detailed insights into this kind of data is necessary. Therefore, researchers need access to intuitive visualization tools to obtain a global overview of the data. Commonly used tools include Krona (Ondov et al. 2011), MEta Genome ANalyzer (MEGAN) (Huson et al. 2007), iTOL (Letunic & Bork 2016) and VAMPS (Huse et al. 2014). These tools can visualize the taxonomic composition of their datasets by exploring the abundances on a species-level or on a broader taxonomic category whereas in case of iTOL, a phylogenetic clustering approach is used. For single organisms, the Integrative Genome Viewer (IGV) (Robinson et al. 2011), Circos (Krzywinski et al. 2009) and Tablet (Milne et al. 2016) are commonly used tools.

In contrast to pie chart visualizations, as offered in Krona, and to bar chart plots as integrated in MEGAN, Sankey diagrams are a good alternative to visualize gene expression data or microbial community compositions over time. These diagrams can indicate the increase or the decrease of data elements in two or more time points and therefore describe the transition of bacterial growth over time. Sankey diagrams are also commonly used in other research areas to highlight changes over time, e.g. in eye dynamics (Burch et al. 2013), medical records (Huang et al. 2015), energy flows in cities (Chen & Chen 2017), energy efficiency (Dietmair & Verl 2009) and voter transition (Fieldhouse & Prosser 2016).

As the costs for sequencing data decreases, more time series data are produced requiring intuitive methods for the visualization of the respective data and especially for researchers that have no direct knowledge or access to programming languages. For the analysis of genes, Sankey plots are an ideal visualization method to detect candidate genes with a similar expression profile. Especially, when interlinked with functional descriptions, such as Interpro (Mulder et al. 2002), gene ontology categories (GO, (Ashburner et al. 2000)), conserved orthologous groups (COG, (Tatusov et al. 2000)), protein families (PFAM, (Finn et al. 2014)) or KEGG (Kanehisa & Goto 2000) to explore the biological context of the genes, these types of visualization provide the basis to obtain a global overview. In our tool, *BioSankey*, we used Sankey plots to analyze microbial communities, both on species and taxonomic level to inspect gene expression along different time points. Furthermore, we can analyze lists of differentially expressed genes (DEGs) by inspecting the expression transitions over time and also functionally by offering customized queries on the data. Additionally, the tool can be easily embedded into an existing analysis workflow. With *BioSankey*, we provide a tool for functional queries to search for and visualize key genes, with an additional export functions to allow the integration of plots into publications.

## Material and Methods

### Import of data into *BioSankey* and generation of the HTML site

The absolute or relative abundance of an element type (e.g. abundance per population, or gene) is provided by the user as a text file or Microsoft Excel file, where a row depicts the data element and the column a time point. There is an additional second column where, optionally, the description of the gene or OTUs can be specified. These identifiers can be then used to query the dataset. Time series data of the fluctuation of up- and down-regulated DEGs can be also provided by the user as multiple files covering the pairwise comparisons between consecutive time points (e.g. first versus second time point), which we will describe in the Use case 1. The whole functionality of *BioSankey* is summarized in Figure 1 starting from the specification of the possible input files, the integration of the information into a HTML site with help of a customized python script and finally the generation of the HTML site and PDF exports.

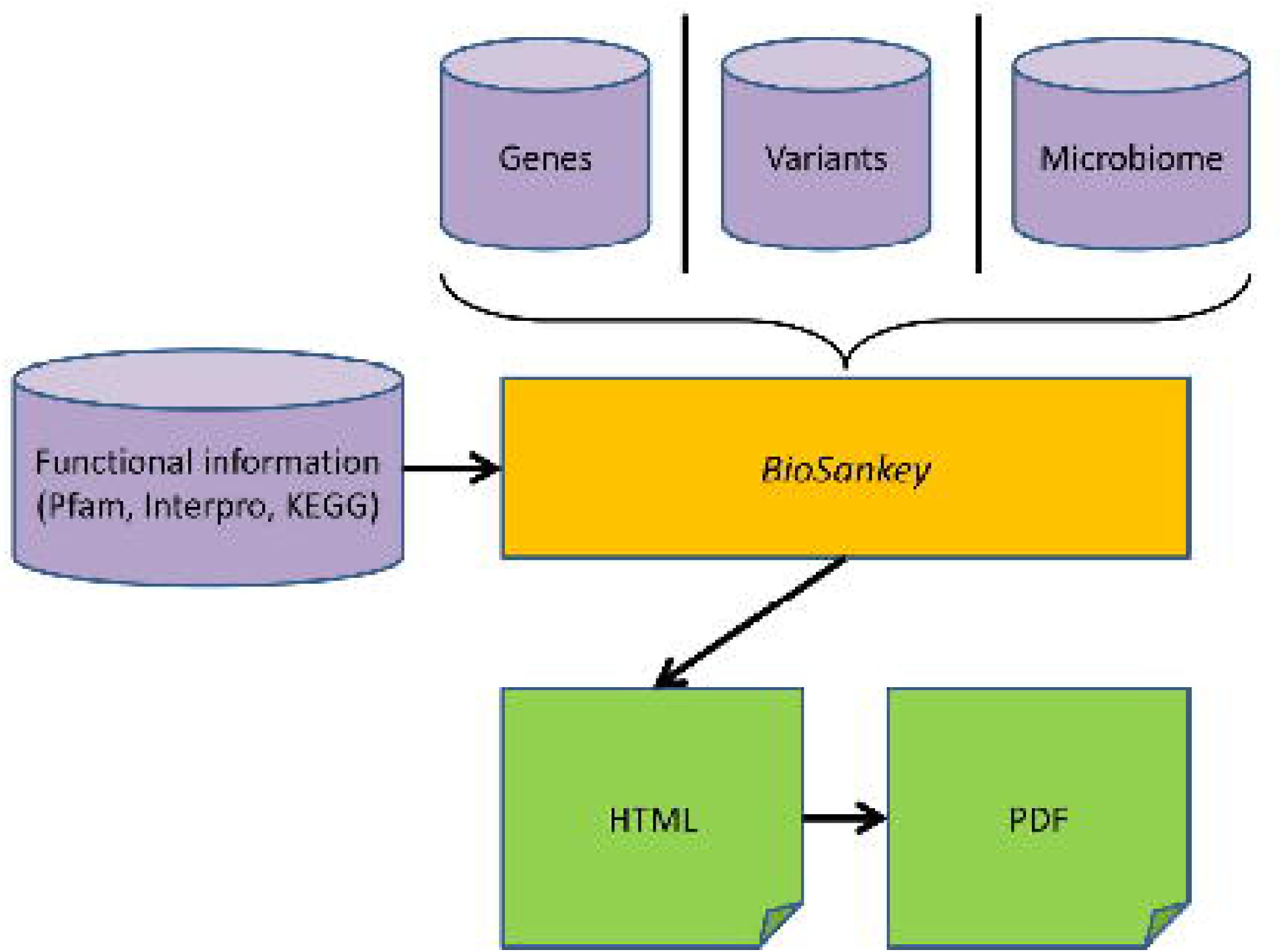

We use one central script to extract the input data such as the abundancy data and functional annotations and optionally DEG lists to generate project-specific HTML pages containing Sankey diagrams based on the Google API. The HTML page includes features to query genes or bacterial species if this data is provided. By using Javascript, this allows to generate visualizations directly without requiring a webserver in the background. Each generated HTML plot can be then exported as a PDF, either silently at the command line or afterwards interactively adjusted.

### Integration of functional information for genes and species

A user can integrate functional information for genes by adding a mapping table during the generation of the website, which allows to search for domain descriptions in the search panel. Thereby, it is possible to search e.g. for a Pfam, GO identifier or for any other integrated descriptive information. In case, that a user provides abundancies of taxonomic species it is possible to select broader taxonomic categories within the visualizations just by entering e.g. a phylum or species identifier. Then, the assignment to taxonomic groups has to be provided in the second column of the abundancy matrix. To this end, a user must provide the taxonomic unit for each operational taxonomic unit (OTU). Instead of a gene identifier, functional information, such as the overlapping gene identifier can be used. Also, in case that differentially expressed genes between different time points were integrated, the tool allows to select those, that are differentially expressed between all timepoints or between selected pairwise time points (e.g. 6h and 12h), supposed this information is provided by the user.

### Visualization of elements in the HTML site

The HTML file contains three different panel views. The first panel is the input box (Supplemental Figure 1), where a user can provide genes of interest (separated by a comma) or can select a functional domain. If no domain information was provided, no selection box is visible. The second panel contains the genes that were previously selected by the search queries. If up to ten genes were selected by the filter criteria, all of them are highlighted as Sankey diagrams, whereas if more than ten genes are in the query set, genes must be selected manually from a selection box, that is embedded in the website. This should provide a good overview of the abundance changes. The third panel contains three visualization modes: A user can select ‘DEG categories’, where an overview of transitions of up- and down-regulated genes between consecutive time points is given, whereas in the mode ‘GENES’ all genes or shown and can be selected. In the ‘METAGENOME’ option, microbial communities are visualized from the highest taxonomic unit to the lowest taxonomic unit, a feature which is inspired by the tools such as Krona and MEGAN to allow to analyse the abundancies of each bacteria and to visualize the abundancy over time. Furthermore, with the input box also selected bacteria can be chosen leading to the update of the ‘METAGENOME’ mode.

## Results and Discussion

### General use of *BioSankey*

*BioSankey* can visualize input data from two different data sources: i) gene expression data, or ii) microbial data. In general, the tool can also be used for any other dataset that contains quantitative data from different time points. When integrated into *BioSankey*, it is possible to infer the fluctuations in abundances in microbial species originating from metagenome projects. These different input data can be then used as input for generating a project-specific HTML site by making use of JavaScript to enable an intuitive and interactive selection and visualisation of all data elements, but also of particular filter criteria by functional description or selection of genes of interest. When the HTML site is generated, the project page, which contains the entire project with all expression information can be exchanged with collaboration partners, as no webserver or database is required. Figure 1 provides the workflow of the tool where two different datasets can be integrated and are then summarized with customized python scripts to generate a HTML page. When the HTML page is generated, we offer an export function for the tool to generate Sankey diagrams in the PDF format.

### Comparability to other tools

We have compared *BioSankey* also to other tools, which also allow to visualize the abundances of species or taxa such as Krona and iTOL. An overview of the functionality that *BioSankey* provides in contrast to the other tools is given in Table 1. Whereas *BioSankey* allows to visualize taxonomical visualisation and time-series data, Krona and iTOL don’t allow to visualize time-series data, but offer a broad range of export functionalities. In contrast to the other tools, *BioSankey* however allows also to visualize the expression of single genes or selected species, which is so far not supported in the other tools.

**Table 1.** Features, that are provided in the three tools *BioSankey*, Krona and iTOL.

| Feature | BioSankey | Krona charts | iTOL |
| --- | --- | --- | --- |
| Taxonomical visualization | Yes | Yes | Yes |
| Time-series data visualisation | Yes | No | No |
| Various Export possibilities | No | Yes | Yes |
| Highlighting of selected genes (e.g. DEGs) | No | Yes | Yes |
| Search function to find genes/taxa of interest | Yes | No | Yes |

### Differential gene expression visualizations

To demonstrate *BioSankey*, we used published gene expression time series data to describe the effects of Camptothecin in U87-MG cell lines to illustrate our feature for visualizing up- and down-regulated genes over time. Camptothecin is a drug, that specifically targets topoisomerase I (Topo I). In their study, authors reported time-related changes and cell line specific changes of gene expression after Camptothecin treatment by considering expression data of six time points (2, 6, 16, 24, 48 and 72h). As a general overview of differentially expressed genes (mode: ‘DEG’) we provide a feature in *BioSankey* to highlight the amount of differential expressed genes as shown in Figure 2. The genes at each time point are filtered for up- and down-regulated expression (in the example a minimal fold-change of 2). In Figure 2 all transitions of genes between these states (up-, down- or low-regulated) in each time point are shown. For a detailed analysis, the genes of a transition (e.g. all genes up-regulated at 2h and at 6h) can be selected and underlying genes can be then visualized separately. To allow this overview visualization, a user must provide lists of differentially expressed genes for each time point in a directory. From the visualization we can observe, that only a certain fraction of the genes (64 of 408) are up-regulated already at the first hours and only 41 of them are then also differentially expressed at the later time points. Furthermore, a user can then select a particular time point of genes that are up-, down- or not differentially expressed and can extract then the respective genes and visualize their expression to find a candidate gene or to obtain a general overview of the functionality of these genes.

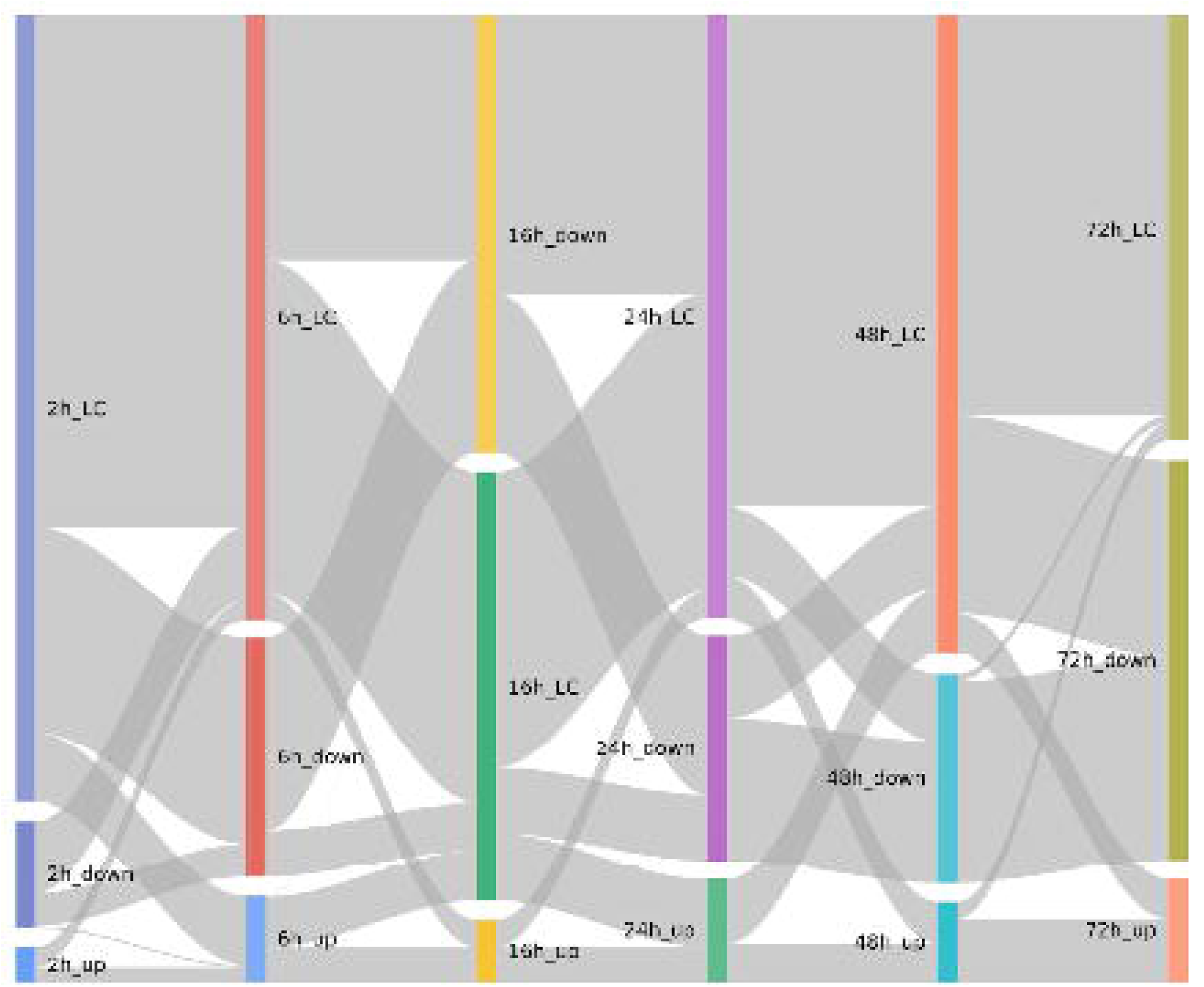

### Microbial community analysis

When metagenomic studies are considered, reads are often assembled with tools such as SSpace (Boetzer et al. 2011) and then used to enter a binning approach such as Maxbin (Wu et al. 2016) and Concoct (Alneberg et al. 2014), which group contigs into Bins based on their tetranucleotide frequencies and of other measurements. As an alternative, more often the sequencing of the 16S rRNA gene is done to assess the abundancy of the species in a metagenomics dataset. Thereby, sequences are clustered based on 97% sequence identity to form OTUs, which can be then analysed with resources such as the SILVA server (Quast et al. 2013) after being processed with mothur (Schloss et al. 2009) and QIIME (Caporaso et al. 2010). In order to demonstrate *BioSankey* for analyzing microbial communities, we have used data from (Caporaso et al. 2011) which contains data generated with QIIME comprising various microbial communities from different tissues. For demonstration purpose, we selected the tongue tissue and extracted all information on genus level, that were specified there and used *BioSankey* to generate a project-specific HTML site. Out of 373 genera, 250 of them had support by at least one read and was integrated into *BioSankey*. We used *BioSankey* to visualize the flow of all genera altogether and used the information to illustrate the abundance of the bacteria in all taxonomical units from bacteria to the final strain and provided by *BioSankey* and selected then two examples (Figure 3).

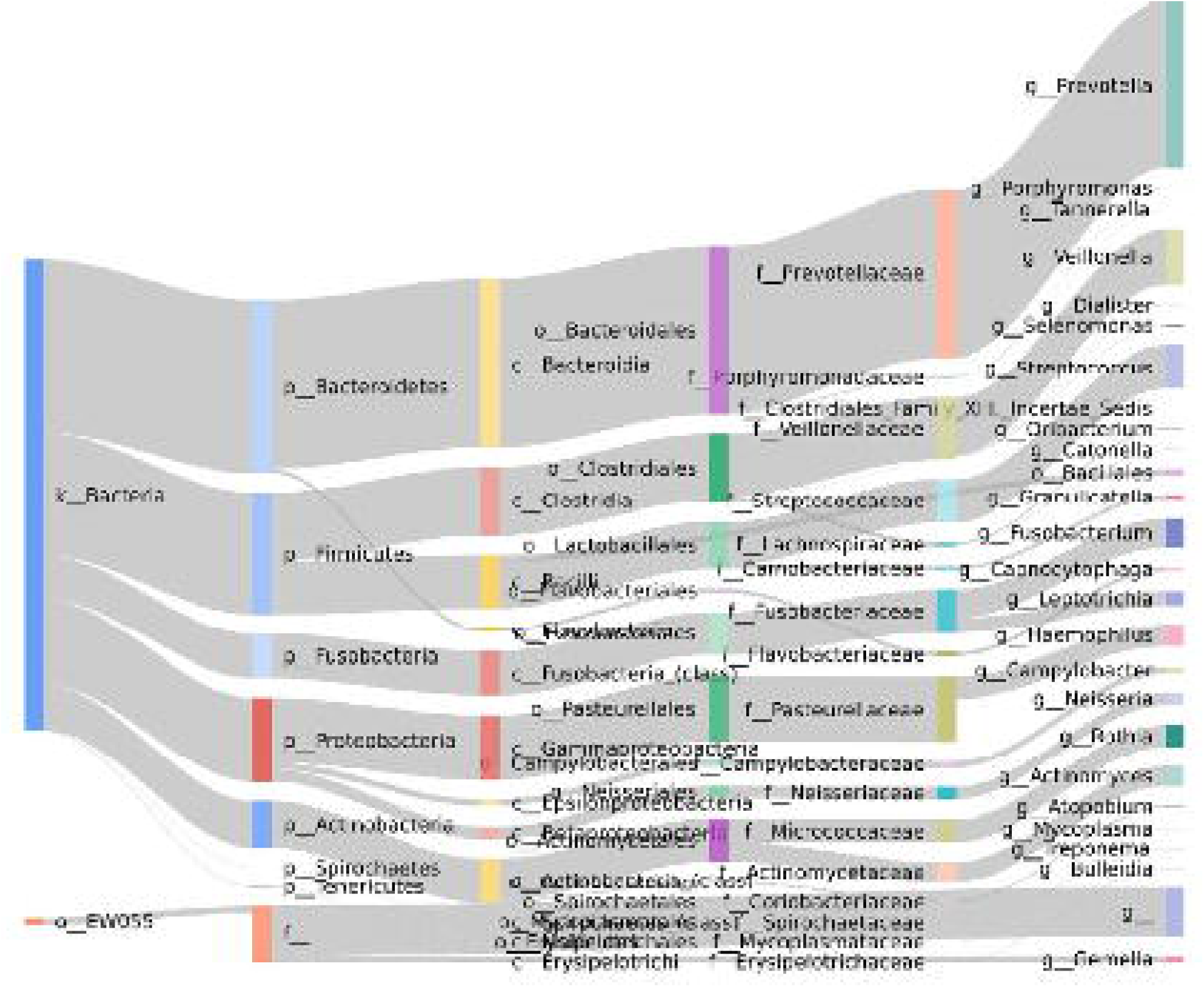

## Conclusion

We have established *BioSankey* as a tool, which offers an alternative way to analyse gene expression or the abundances of metagenomics datasets over time by using Sankey diagrams, functional enrichment analysis and overview panels without the requirement of dependences of web servers and databases. We demonstrate the possibilities to interactively view the data for an efficient analysis. *Biosankey* is a valuable tool to get insights and understand the complexity of different datasets, from a high-level view of gene numbers to the genes, and in case of metagenomics, from a taxonomic high level down to strain level. This tool is important for researchers, who want to analyse the taxonomic composition of bacterial species in metagenomes by *e.g.* also selecting broader taxonomic categories. As an additional feature of the tool data exchange with collaborations is easily accomplished. For the dual RNA-seq experiment, *BioSankey* might be especially powerful when hosts and symbionts are compared to each other or in order to efficiently detect interesting candidate genes. Therefore, we have integrated various criteria, which are based on domain functionalities or gene description information.

## Material availability

The software is available at the Github repository https://github.com/nthomasCUBE/BioSankey.

